# High-accuracy protein model quality assessment using attention graph neural networks

**DOI:** 10.1101/2022.09.24.509136

**Authors:** Peidong Zhang, Chunqiu Xia, Hong-Bin Shen

## Abstract

Great improvement has been brought to protein tertiary structure prediction through deep learning. It is important but very challenging to accurately rank and score decoy structures predicted by different models. CASP14 results show that existing quality assessment (QA) approaches lag behind the development of protein structure prediction methods, where almost all existing QA models degrade in accuracy when the target is a decoy of high quality. How to give an accurate assessment to high-accuracy decoys is particularly useful with the available of accurate structure prediction methods. Here we propose a fast and effective single-model QA method, QATEN, which can evaluate decoys only by their topological characteristics and atomic types. Our model uses graph neural networks and attention mechanisms to evaluate global and amino acid level scores, and uses specific loss functions to constrain the network to focus more on high-precision decoys and high-precision protein domains. On the CASP14 evaluation decoys, QATEN performs better than other QA models under all correlation coefficients when targeting average *LDDT*. QATEN shows promising performance when considering only high-accuracy decoys. Compared to the embedded evaluation modules of predicted *C*_*α*_-*RMSD* (*pRMSD*) in RosettaFold and predicted *LDDT* (*pLDDT*) in AlphaFold2, QATEN is complementary and capable of achieving better evaluation on some decoy structures generated by AlphaFold2 and RosettaFold themselves. These results suggest that the new QATEN approach can be used as a reliable independent assessment algorithm for high-accuracy protein structure decoys.

## 1 Introduction

Proteins are involved in almost all biological processes and cellular functions. Functions of proteins highly depend on their spatial structures, namely the three-dimensional arrangement of amino acids [1]. Experimental tests of protein folding are generally expensive and time-consuming [2]. With the development of high-throughput sequencing, protein sequence data accumulate rapidly [3]. In order to bridge the gap between protein sequence and structure database, the development of in silico tertiary structure prediction methods has been a long-term challenging topic in structural bioinformatics [4-13]. The Critical Assessment of Techniques for Protein Structure Prediction (CASP) is designed to benchmark the progress of protein structure prediction methods. In 2020, The latest artificial intelligence algorithm, AlphaFold2 [6] and RosettaFold [14] have revolutionized this field with performance in CASP 14. Their algorithms have the capability of making accurate protein structure predictions comparable to the lab experiments-based solution. However, there is still a lack of independent convincing computational methods for ranking high-accuracy decoys.

Quality assessment (QA) aims to estimate the accuracy of a protein decoy in reference to its native structure, but without the knowledge of its ground truth [15]. Global distance test-total score (*GDT-TS*) [16] and local distance difference test (*LDDT*) [17] are widely used to measure the difference between the predicted and native structures. As described in the CASP13, significant advances in QA methodology development have been witnessed [18].

Current QA techniques can be generally divided into two categories: single-model methods which operate on a single protein model to estimate its quality, and multi-model methods which use consistency between several candidates to estimate their qualities [19]. Multi-model methods typically have a pool of decoys that are generated by different methods or from different templates, and assume that correct fragments of the target structure are already embedded in the pool. Hence, they evaluate the accuracy of the current decoy by clustering and extracting the consensus information in the pool. Obviously, the performance of multi-model methods depends on the quality and size of the pool [20], and an excellent multi-model QA approach often needs a large pool of dozens of models and hundreds of decoys, resulting in high computational cost accordingly [21]. In contrast, single-model methods usually extract inherent features without external predictors. They mainly are based on the topology and energetic analysis of a single decoy and predict the distance from the current conformation to the lowest energy conformation through structural analysis. Multi-model methods achieve high performance in CASP13 due to the significant improvement of structure prediction methods [19]. Since the recent protein structure methods often have already integrated multiple models, the gap between single-model and multi-model QA methods is becoming narrowed.

The single-model methods are closer to indicate the distance from current conformation to the lowest energy conformation and have attracted increasing attention in CASP competition. In the recent CASP14, single-model QA methods account for more than 70% of all model quality assessment methods. They typically use different feature combinations and machine learning methods to learn the implicit relationship between decoy structure and its quality. For instance, in ProQ2 and ProQ3 [22, 23], features such as atom-atom contacts, residue-residue contacts, surface exposure, predicted and observed secondary structures, and Rosetta energy terms are fed into support vector machines (SVMs) for calculation.

With the development of deep learning techniques, more neural network-based (NN-based) methods are proposed. ProQ3D [24] replaces the SVM in ProQ3 with a multi-layer perceptron model, and ProQ4 uses more protein structural features such as dihedral angles, secondary structures, hydrogen bond energy, and co-evolutionary information for training with multi-stream structure. DeepQA [25] uses a deep belief network to make predictions by using structural, chemical, and knowledge-based energy scores. A more intuitive idea in recent years is to feed decoy structures into 3D convolutional networks or graph neural networks. 3DCNN_MQA [26] and Ornate [27] project the decoy structure onto a 3D grid and use 3D convolution to convert this voxelized representation into a quality score. On the other hand, ProteinGCN [28] and GraphQA [29] represent decoy structures on the graph at the atomic level and amino acid level, respectively, and then automatically learn the best embeddings from the original node and edge features for local and full-graph attribute prediction. ModFOLD7 [30] and ModFOLD8 [31] are consensus approaches, which integrate single-model methods such as CDA, SSA, ProQ, ProQ2D, ProQ3D, VoroMQA, DBA, MF5s, MFcQ and ResQ7 [32]. This series of methods all top the CASP13 and CASP14 competitions.

However, according to CASP14, the QA is lagging behind the development of protein structure prediction methods [33], where existing QA algorithms could not provide accurate assessment for high-accuracy decoys. Although the overall performance of these QA models in CASP14 is acceptable, for instance, they can still distinguish the poor decoys (*GDT-TS*<0.5), but when considering those decoys close to native structures (e.g., *GDT-TS*≥0.5), a significant drop in QA accuracy is observed. This highlights the importance of developing protein quality assessment (QA) for high-accuracy models with the available of some excellent structure prediction algorithms, such as AlphaFold2 and RosettaFold. The potential reasons for the failure of existing QA methods on the high-accuracy decoys are the features used in QA model, such as homology and coevolution information, which have already been widely used and learned in different structure prediction models’ networks. This feature overlap may confuse the QA model when evaluating outputs from different structure predictors. A proper estimation of the free energy differences between decoys instead of trying to mine more homology or coevolution features is potentially one of the keys to improving QA algorithm.

In this study, we propose QATEN, a novel single-model quality assessment method based on graph neural networks and the attention mechanism. Protein coevolutionary information which was widely used in structure prediction is not required by QATEN. It only considers types of heavy atoms in amino acid and topology information as the input. We also design a novel loss function to enhance the attention to high-accuracy decoys and more realistic local segments in one decoy. QATEN has much fewer trainable parameters and faster speed than another computationally efficient model Ornate [27]. On the CASP14 datasets, QATEN is highly competitive compared with all other single-model QA methods, and achieves state-of-the-art performance on high-precision samples. On some decoys predicted by AlphaFold2 and RosettaFold, our model can provide more correlated predictions.

## 2 Datasets

Nine benchmark datasets are used in this study. DeepAccNet dataset and GNNRefine dataset are merged for training and validation [33, 34], and we randomly select 5% of the decoys from the two datasets to form the validation dataset. Supplementary Fig.S1 shows the decoy quality distribution of these two datasets. The other seven datasets are used as independent test sets. Details of the datasets are provided below.

1. DeepAccNet dataset [34]: this dataset contains 1,104,080 decoys generated from 7,992 protein sequences. The proteins are retrieved from the PISCES server and deposited to PDB by May 1^st^, 2018 and ranging from 50 to 300 residues in length and having resolutions less than 2.5 Å. Decoys in this dataset are generated using three methods: comparative modeling, native structure perturbation, and deep learning guided folding. All decoys are subject to dual-space relaxation in Rosetta to mitigate the possible difference in modeling procedures between different methods.
2. GNNRefine dataset [35]: this dataset contains 509,443 decoys generated from 29,455 protein sequences. The protein chains are selected from CASP7-12 and CATH domains (sequence identity <35%) released in March 2018. Decoys are modeled by 13 template-based and template-free models for each protein using their protein structure prediction software RaptorX. Compared with DeepAccNet set, this set has fewer protein chains and more decoys for each protein, and decoys have lower average quality.
3. CASP14: this dataset includes 3897 decoys corresponding to 26 target proteins (about 150 per target protein) provided by the CASP14 experiment. These decoys are submitted by participating servers and screened by the organizers for EMA (model accuracy evaluation) experiments. To guarantee at least 90% sequence structural integrity, we abandon eight protein chains with many missing coordinates in the experimental observations proposed by the organizers. Similar to DeepAccNet dataset, all decoy structures are dual-space relaxed in Rosetta software. A list of these 26 target proteins can be found in Supplementary Table.S1.
4. CASP14_GDT & CASP14_LDDT: this dataset is composed of decoys with *GDT-TS*≥0.5 and the decoys with average *LDDT*≥0.5, which are considered as the high-accuracy decoys, in the CASP14 dataset. They have a total of 1193 decoys and 1362 decoys respectively.
5. CASP14_alphafold & CASP14_rosettafold: as the most advanced models in the structure prediction field, AlphaFold2 provides predicted *LDDT*, and RosettaFold provides predicted *C*_*α*_**-** *RMSD* to assess their 5 final predictions at the amino acid level. Thus, we successfully use AlphaFold2 to predict 130 decoys and RosettaFold to predict 120 decoys in CASP14 (we did not get predictions for two proteins from RosettaFold), and the names of proteins are shown in Supplementary Table.S1.
6. RCSB_alphafold & RCSB_rosettafold: 250 experimentally solved protein structures with high accuracy (resolution less than 1.2Å) were downloaded from RCSB PDB (https://www.rcsb.org/) where they were deposited from July 2021 to May 2022. In this dataset, each protein has a complete experimental structure derived from X-ray diffraction, and the complete chain A is isolated for evaluation. The length of these protein sequences ranges from 20 to 800, and most proteins contain 80 to 400 amino acids. Compared with CASP14 dataset, protein chains in this set contain significantly more homologous information, which is beneficial to the structure prediction models but not conducive to the QA models. The sequence of amino acid residues in the protein structure was fed into AlphaFold2 and RosettaFold to predict five decoys respectively. Accordingly, RCSB_alphafold has 1250 decoys and RCSB_rosettafold has 1230 decoys (RosettaFold failed to predict four protein chains). The names of all proteins are given in Supplementary Table.S1.

## 3 Results

### 3.1 Overview of QATEN pipeline

The QATEN is the first GNNs-based model focusing on the ranking of high-precision decoy structures, to the best of our knowledge. It uses the attention mechanism to mine structural features and designs loss function to constraint model to pay more attention to high-precision decoys and high-precision domain of decoys. Fig.1a shows the flowchart of QATEN which mainly includes three steps: 1) extract atomic and geometric features from the initial structure and represent the initial decoy structure as a graph, 2) predict and update the node embeddings in the graph using graph neural networks (GNNs), and 3) calculate the graph score using graph pooling at the global protein level as well as at the local residue level.

**Fig.1.**
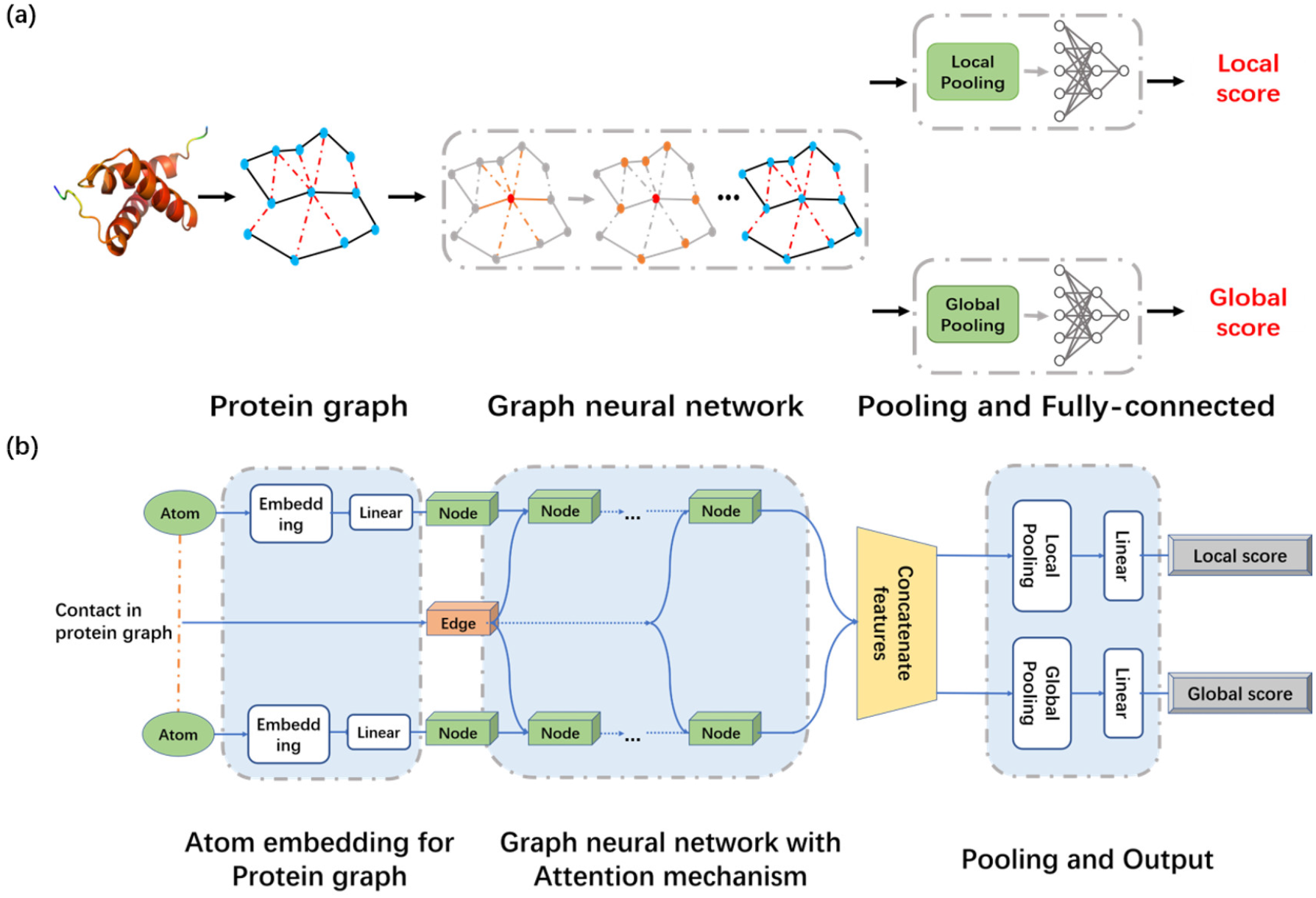
(a) The flowchart of QATEN which mainly includes three steps: 1) extract atomic and geometric features from the initial structure and represent the initial decoy structure as a protein graph, 2) predict and update the node embeddings in the graph using GNNs, and 3) calculate the result scores through different pooling network. (b) The network architecture of QATEN. The original features of a pair of connected atoms in the protein graph are represented as node features and edge features by embedding layers. Edge features remain unchanged and are spliced at corresponding nodes, and node features are updated in GNNs. Node features are concatenated and expanded as one dimension, then sent into downstream pooling networks.

The GNN-based embedding prediction is the key factor affecting the QA model quality. The graph neural networks in QATEN mainly consists of five layers of modules with the same network architecture. Each module would get protein graph representations with node features and edge features learned in the last module. The protein graph is built on the atom feature (node) and distance or contact feature (edge) between the 50 closest atom pairs in structure (detailed in the Methods section). By going through multi-head attention layers, the node features are updated to capture structural information. As shown in Fig.1b, the original features of a pair of connected atoms in the protein graph are represented as node features and edge features by embedding layers. When passing through each module of GNNs, edge features remain unchanged and are spliced at corresponding nodes. The node features after GNN learning and updating are expanded as one dimension and the downstream results were obtained by two pooling networks.

### 3.2 Evaluation metrics

For the main experiments, we restrict the measures of global performances to *GDT-TS* and the average *LDDT*, since they are widely used and the official scores in the CASP competition. For each QA method, we consider the predicted and ground-truth scores to compute Pearson (𝒫), Kendall (𝒦), and Spearman (𝒮) correlation coefficients across all decoys of all targets. Detailed explanations of the three correlation indicators are available in supplementary information. We define decoy label with *GDT-TS* or average *LDDT* less than 0.5 as 0, vice versa as 1, and calculate AUC defined as the area under the ROC curve. In addition, we also compare the time cost of different methods.

Particularly, RosettaFold and AlphaFold2 also provide an assessment of amino acid residue levels for their predicted decoy structures. AlphaFold2 predicts *LDDT* in its evaluation module while RosettaFold predicts *C*_*α*_-*RMSD*. For these two models, we calculate Pearson coefficients between *pLDDT* and *LDDT, pRMSD*, and *RMSD* on their own predicted decoys respectively.

### 3.3 Performance on the validation dataset and CASP14 dataset

Fig.2 shows the correspondence between QATEN predictions and real measures on the validation set and CASP14 test set. The prediction results of all decoy structures of these target proteins not included in the training set are well correlated with *GDT-TS* and average *LDDT*. Almost all of the samples are distributed around the positive correlation line (*y* = *x*). Overall, although the detailed prediction is not pixel perfect (for example, in the CASP14 test set, the predicted value of some decoy structures is lower than the actual *GDT-TS* score), our results still show a good correlation on both indicators and are sufficient to provide good support for the structure prediction model.

**Fig.2.**
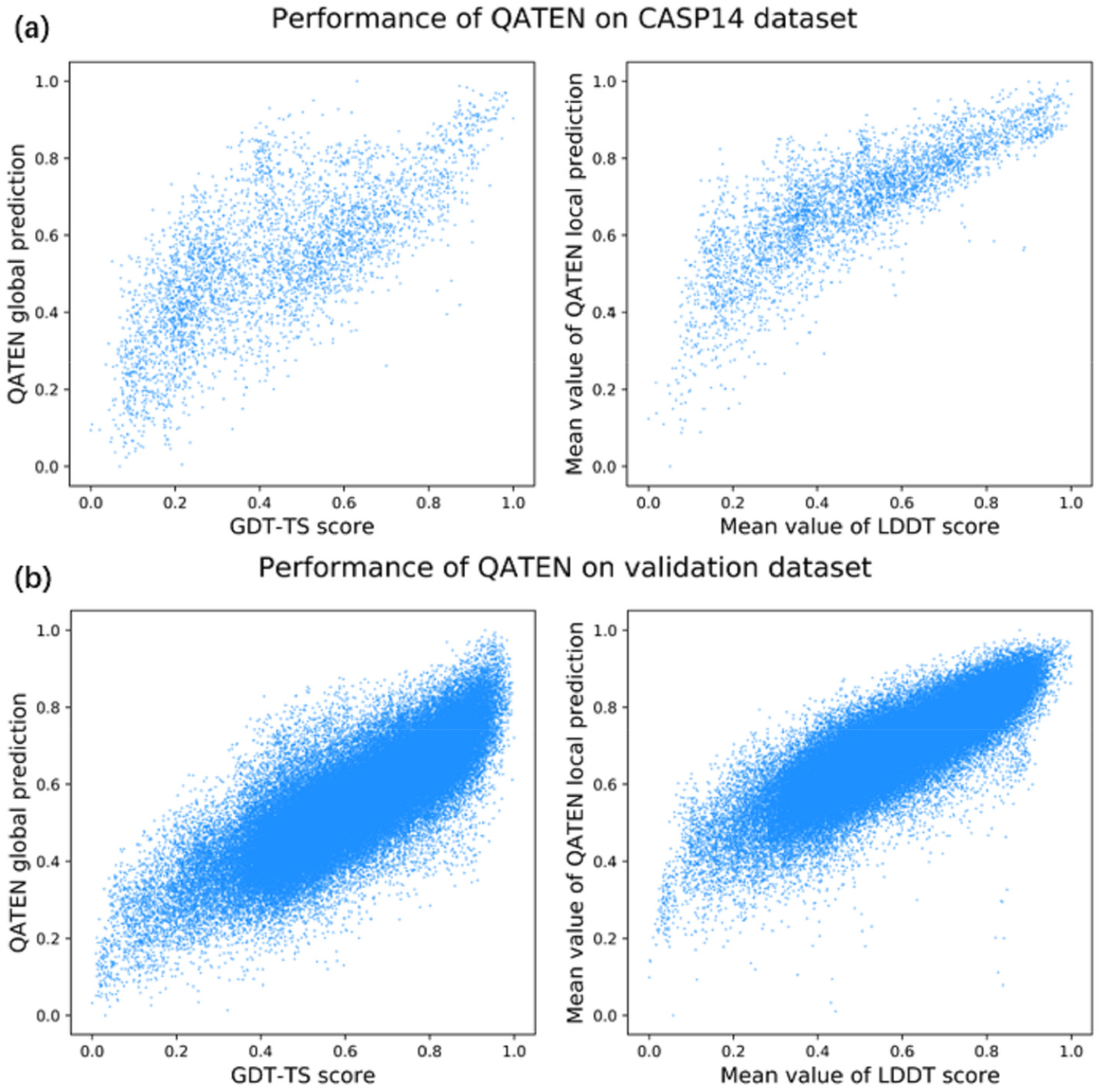
The correspondence between QATEN predictions and real measures on the validation set and CASP14 test set. (a) 𝒫 is 71.77% on CASP14 dataset (3897 decoys) when considering *GDT-TS* as the metric. And 𝒫 is 82.17% on CASP14 dataset when considering the mean value of *LDDT* as the metric. (b) 𝒫 is 81.25% on the validation dataset (87634 decoys) when considering *GDT-TS* as the metric. And 𝒫 is 84.74% on the validation dataset when considering *LDDT* as the metric.

### 3.4 Comparison with other QA models

We test QATEN on the decoys of 26 CASP14 protein targets and compare it with all single-model QA methods in CASP14 [33]. Besides, we also compare QATEN with some publicly available software such as DeepAccNet, DeepAccNet-Bert [34], ProteinGCN [28], and ZoomQA [36]. We directly use their released model parameters. We compare 10 models with the highest score under each criterion here, and put the complete results of 48 baselines in the supplementary information.

Performance of QATEN is slightly lower than ModFold8 and ModFold8_rank, comparable to DeepAccNet while better than the others on all the 𝒫, 𝒦, and 𝒮 correlation coefficients when targeting *GDT-TS* (Fig.3a). As shown in Fig.3b, QATEN performs better than the others on all the 𝒫, 𝒦, and 𝒮 correlation coefficients when targeting average *LDDT*. It is worth mentioning that QATEN only used a much smaller network with about 80k trainable parameters, which is 25 times fewer than other relatively simple networks like Ornate.

**Fig.3.**
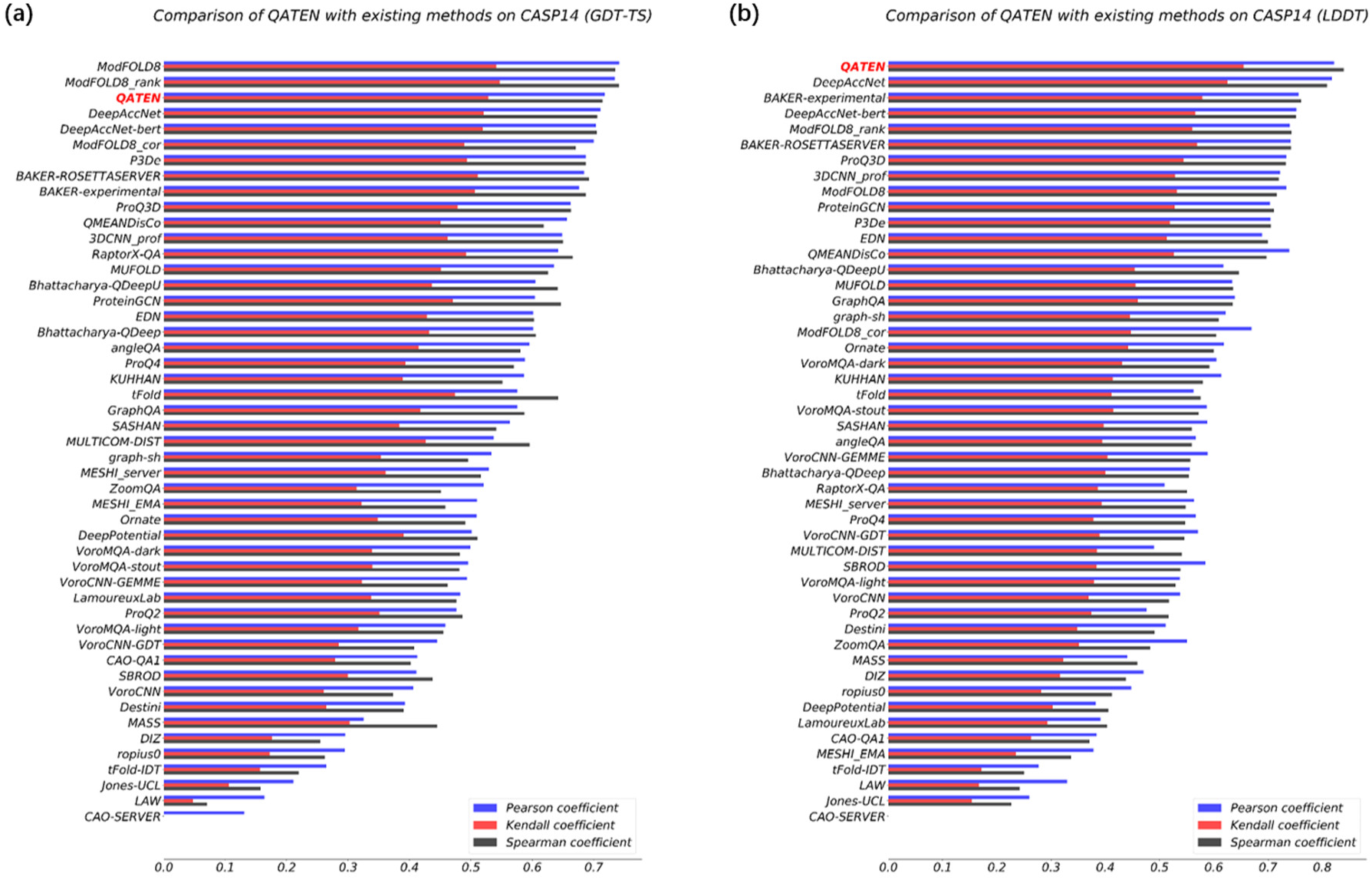
Ranking performance of QATEN and other baseline models on CASP14 dataset by Pearson (𝒫), Kendall (𝒦), and Spearman (𝒮) correlation coefficients. (a) Using *GDT-TS* as the measure. (b) Using average *LDDT* as the measure.

QATEN achieves the best on CASP14_GDT and CASP14_LDDT when considering the correlation coefficient on high-precision decoys (Fig.4a and Fig.4b). When considering high-precision decoys, we observe a significant decrease in accuracy in most models which can be demonstrated by comparing Fig.3 and Fig.4. A potential reason is that QA models focus on learning coevolutionary information, which has been done by structure prediction methods. Traditional QA models tend to separate high-accuracy decoy structures while may neglect ranking and selection of them. In fact, with the success of AlphaFold2 and RosettaFold, as well as advances of other structural prediction models on CASP14, a significant amount of high-precision decoy structures are predicted, which makes QA models underperform on the high-accuracy decoys. Detailed values of the comparison in Fig.3 and Fig.4 are given in Supplementary Table.S2 and Table.S3.

**Fig.4.**
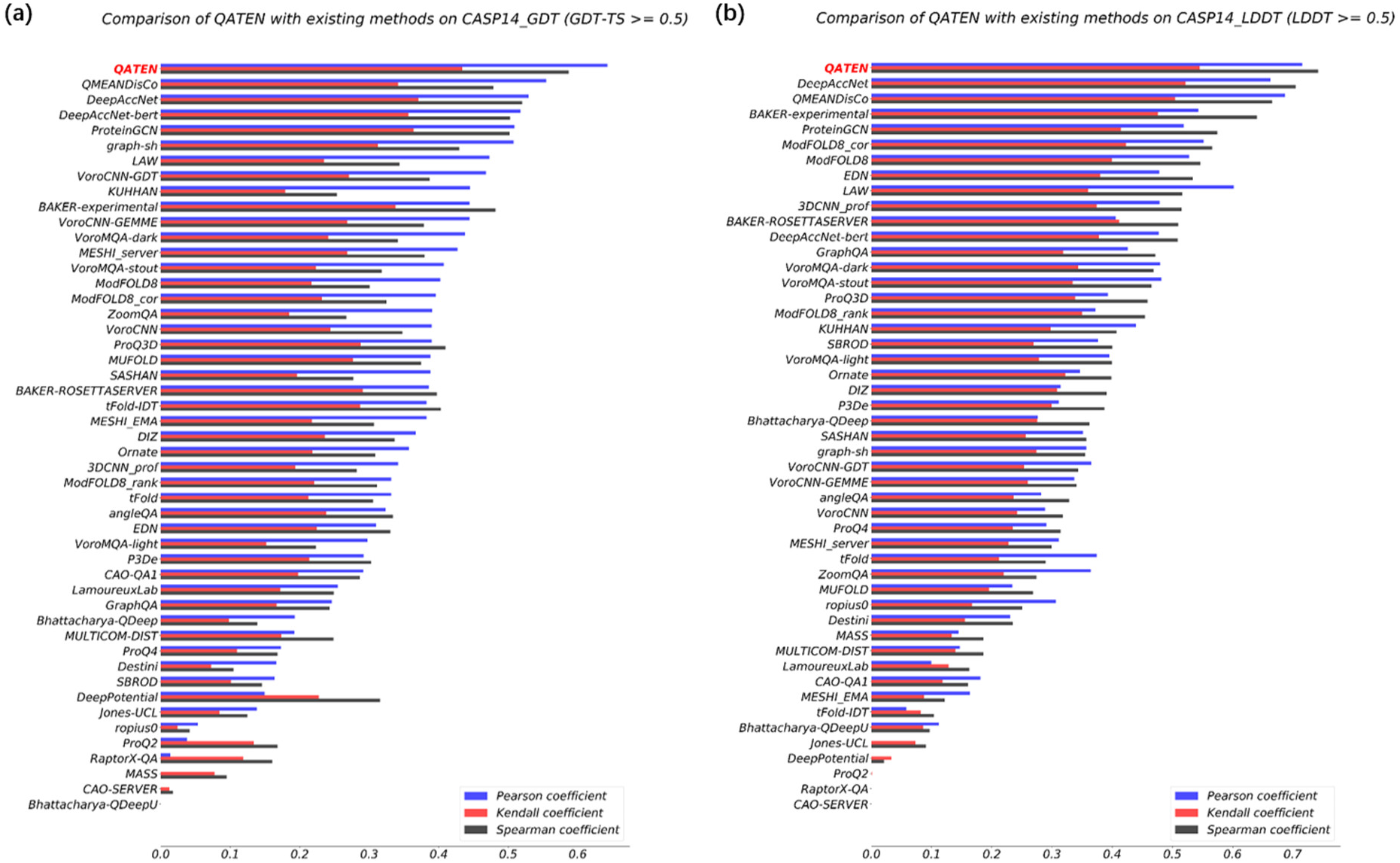
Ranking performance of QATEN and other baseline models on CASP14 dataset by Pearson (𝒫), Kendall (𝒦), and Spearman (𝒮) correlation coefficients. (a) Considering GDT-TS as a measure. (b) Considering average *LDDT* as a measure.

Table 1 shows the results of our further comparison of high-precision decoys, which were further divided into two parts: relatively accurate (0.5≤*GDT-TS*<0.8) and highly accurate (*GDT-TS* ≥ 0.8), and QATEN achieves the best performance on correlation indicators. A complete comparison list could be found in Supplementary Table.S4.

**Table 1.**
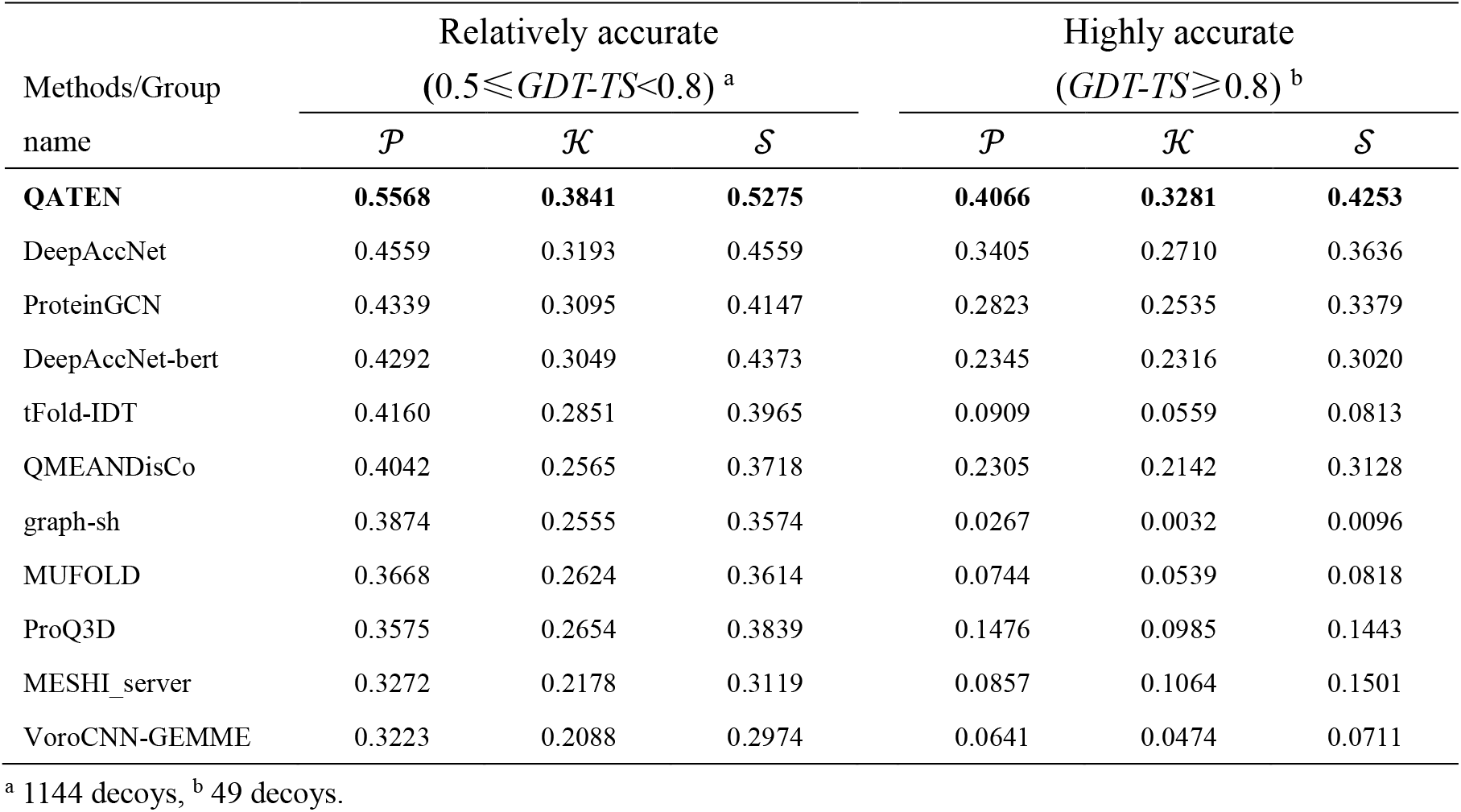
Ranking performance of QATEN and other top-10 baseline models on different bins of accuracy (using *GDT-TS* as the distinguishing criterion).

When calculating AUC, we define decoy label with *GDT-TS* or average *LDDT* less than 0.5 as 0, and vice versa as 1. Since AUC is defined as the area under the ROC curve, it is expected that this indicator reflects the ability of each QA model to distinguish high-accuracy decoys from low-quality decoys. Table 2 shows the comparison of QATEN with baselines on AUC (only the top-10 models are selected). The complete comparison is shown in Supplementary Fig.S2. QATEN performs comparably to DeepAccNet and is better than others when considering average *LDDT*. And it performs only slightly worse than ModFold8 [31], DeepAccNet, and P3De (Human) when considering *GDT-TS*.

**Table 2.** Ranking performance of QATEN and other top-10 baseline models on *AUC*.

| Methods/Group name | *AUC*<sup>a</sup> | *AUC*<sup>b</sup> |
| --- | --- | --- |
| DeepAccNet | **91.68%** | 84.75% |
| **QATEN** | 91.64% | 84.31% |
| BAKER-experimental | 88.37% | 82.57% |
| DeepAccNet-bert | 87.62% | 84.75% |
| BAKER-ROSETTASERVER | 87.26% | 83.09% |
| ProQ3D | 87.26% | 82.83% |
| ModFOLD8_rank | 86.66% | **85.55%** |
| P3De | 86.20% | 84.86% |
| END | 86.04% | 80.04% |
| ModFOLD8 | 86.00% | 85.08% |
| QMEANDisCo | 85.65% | 79.47% |

*AUC*<sup>a</sup> set mean value of *LDDT* > 0.5 as label 1; *AUC*<sup>b</sup> set *GDT-TS* > 0.5 as label 1.

### 3.5 Comparison between QATEN and confidence predictor in AlphaFold2

The confidence predictor in AlphaFold2 uses *pLDDT* to make the local evaluation of its predicted decoy structures. We compare QATEN and confidence predictor in AlphaFold2 separately on CASP14_alphafold and RCSB_alphafold datasets. For 130 decoys in CASP14_alphafold, the average 𝒫 of local assessment is 0.6368 by QATEN and 0.647 by confidence predictor in AlphaFold2. For 1250 decoys in RCSB_alphafold, the average 𝒫 of local assessment is 0.5843 by QATEN and 0.6066 by confidence predictor in AlphaFold2. Protein structures in CASP14_alphafold are simpler than those in RCSB_alphafold, which reduces the difficulty in quality assessment and both QA models perform well. In the end-to-end training of AlphaFold2, the confidence predictor is also involved in the training and recycling process [6]. Protein chains in RCSB_alphafold have more homologous information, consequently helping widen the gap between QATEN and AlphaFold2.

Since we consider a QA model evaluates a decoy well when 𝒫 on this decoy is higher than 0.5. To further compare the two models, we define a hit decoy as 𝒫 by a model on current decoy is higher than 0.5 and a hit protein as 𝒫 by a model on all 5 decoys of current protein are higher than 0.5. Table 3 shows the number of hit decoys and hit proteins by QATEN and confidence predictor in AlphaFold2. It can be seen that QATEN is slightly better than AlphaFold2 in both CASP14_alphafold and RCSB_alphafold datasets. It is worth noting that the training and recycling process makes confidence predictor only available to decoys generated by AlphaFold2, but QATEN can be used on decoys generated by any other different structure models.

**Table 3.** Comparison between QATEN and confidence predictor in AlphaFold2.

| Dataset | Method | $N_{hit_{decoy}}^a$ | $N_{hit_{protein}}^b$ |
| --- | --- | --- | --- |
| RCSB_alphaFold | **QATEN** | **913** | **137** |
|  | *pLDDT* | 873 | 130 |
| CASP14_alphaFold | **QATEN** | **105** | **18** |
|  | *pLDDT* | 102 | 17 |

<sup>a</sup>: hit number of decoys; <sup>b</sup>: hit number of proteins.

We merge CASP14_alphafold and RCSB_alphafold datasets into a new dataset and further explore from the perspectives of multiple sequence alignment (MSA) depth and secondary structure. Fig.5 shows the distribution of hit decoys according to MSA depth (MSA is searched and generated by AlphaFold2). As shown in Fig.5, QATEN performs comparably to AlphaFold2 when MSA is less than 1000 and underperforms than confidence predictor in AlphaFold2 when MSA is higher than 2000. However, QATEN can better evaluate these decoy structures when MSA depth is in the range of 1000-2000. An interesting conjecture is that when the MSA information is rich, AlphaFold2 could predict the structure well and obtain generally high confidence (Alphafold2 hit 93.75% decoys with MSA depth above 3000); when the MSA information is poor, AlphaFold2 would adopt a more cautious prediction and evaluation strategy; and when the MSA information is relatively insufficient, it may overestimate the prediction but ignore some parts with unsatisfactory prediction.

**Fig.5.**
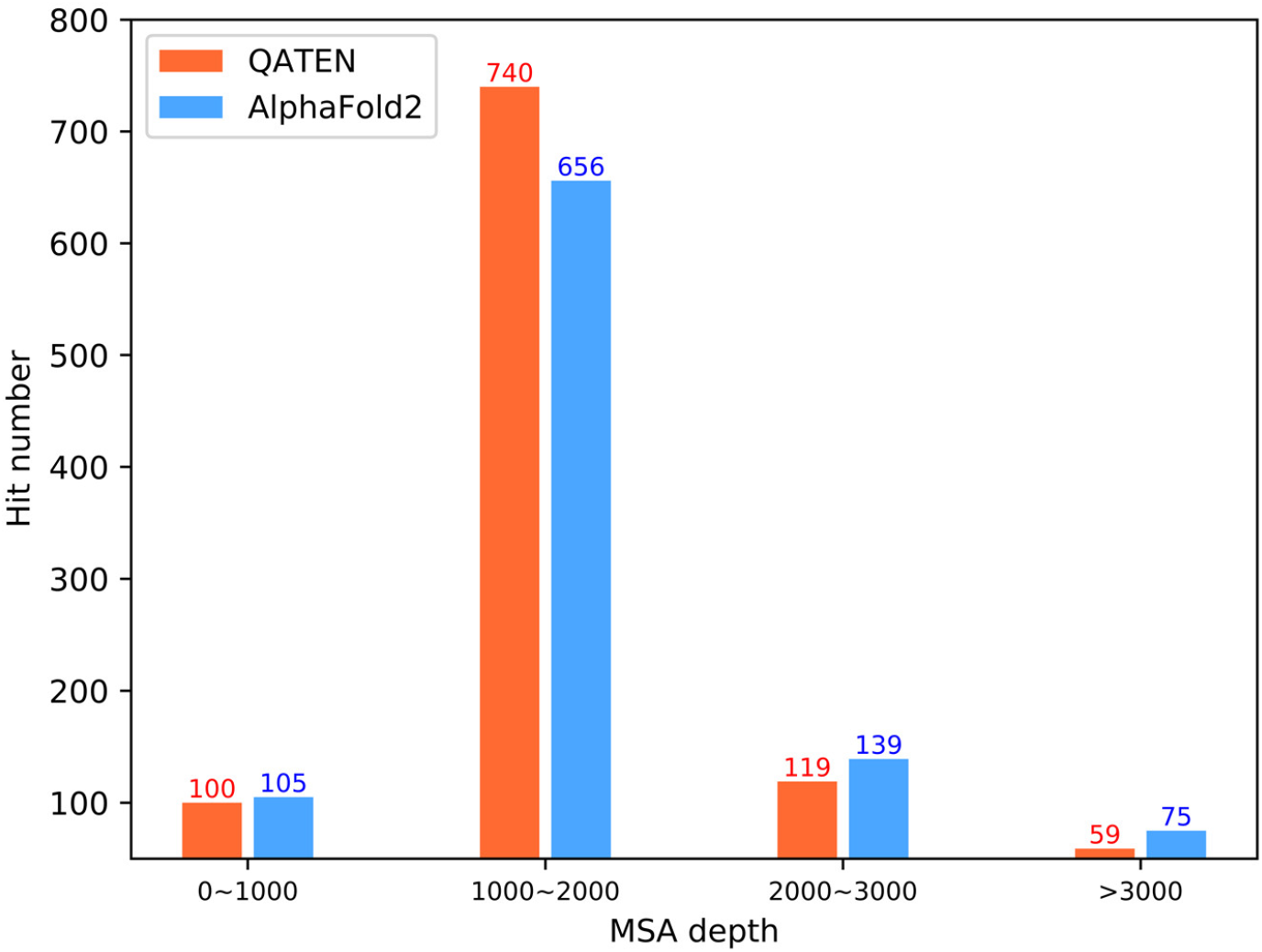
Distribution of the hit decoys according to MSA depth. QATEN performs better on decoys with MSA depth between 1000 and 2000, while the confidence predictor in AlphaFold2 hit almost all decoys (93.75%, 80 decoys in total) with MSA depth above 3000

We find 191 decoys only hit by QATEN and 148 decoys only hit by confidence predictor in AlphaFold2, and count the distribution of secondary structure elements using DSSP [37] software. Fig.6 shows the distributions of secondary structure elements in decoys only hit by (a) QATEN or (b) AlphaFold2. The distribution of secondary structure elements in all decoys is also shown in (c). The distribution of secondary structure elements in two sets mainly concentrated on Strand, *β* bridge and longer sets of hydrogen bonds and *β* bulges, and Helix, *α* helix and *π* helix. Compared to general decoy in CASP14_alphafold and RCSB_alphafold datasets, more strands (+6%) and less helix (−3.2%) could be significantly observed on those decoys only hit by QATEN.

**Fig.6.**
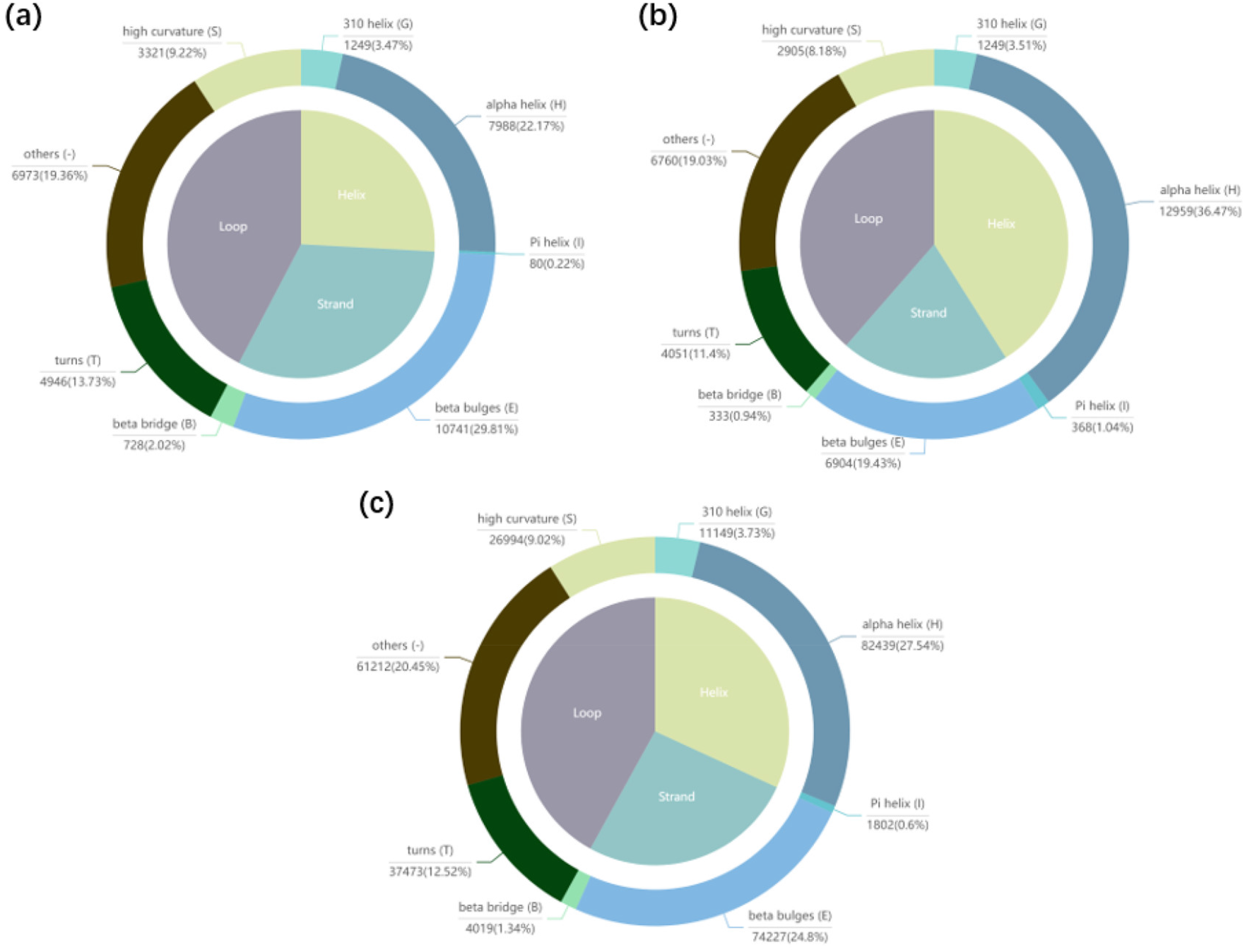
Distribution of secondary structure elements in decoys only hit by (a) QATEN or (b) confidence predictor in AlphaFold2. Additionally, the distribution of secondary structure elements in all decoys is also shown in (c).

But less strand (−5.8%) and more helix (+9.2%) could be observed on those decoys only hit by confidence predictor in AlphaFold2. These results indicate that QATEN could be used as an extra QA tool for the decoys generated by AlphaFold2.

In Fig.7, we make six cases comparison between the QATEN and the confidence predictor in AlphaFold2. Each graph represents a decoy sample predicted by AlphaFold2, and the six graphs represent 5SBV-model-3, 7B5U-model-5, T1031-model-4, T1039-model-1, T1029-model-3 and T1033-model-4 decoys respectively. The first subgraph (gray) represents the native structure. We align the native structure with the decoy structure for better comparison. The second to fourth subgraphs all represent the decoy structure, and the colors on the structure represent the score distribution of different evaluation methods (*LDDT*, score of QATEN, *pLDDT* of AlphaFold2), and the mean value of the score are shown under structures. The range of color (red-orange-yellow-green-cyan-blue-purple) is correlated with the range of scores (1-0).

**Fig.7.**
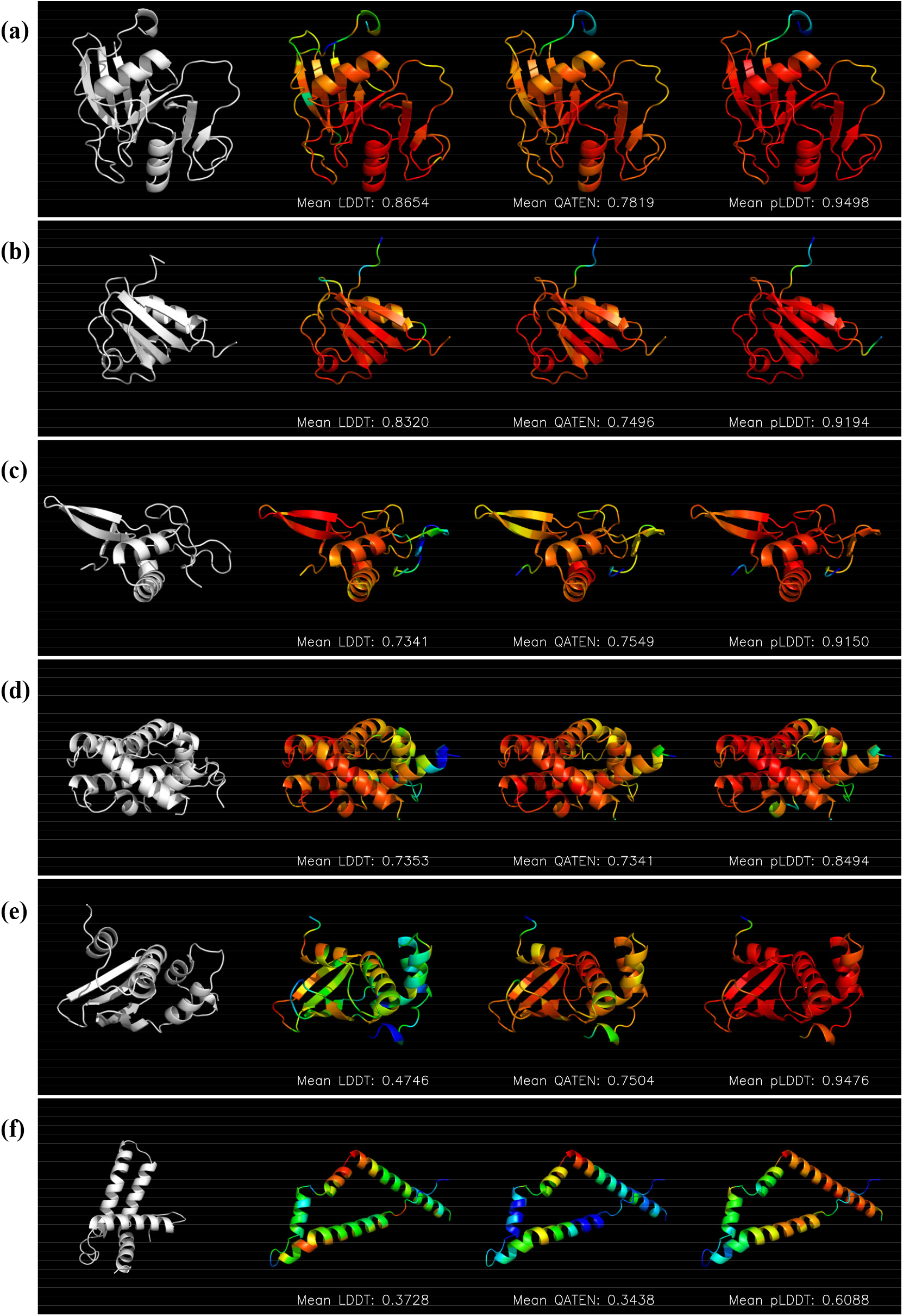
Comparison of QATEN and *pLDDT* of AlphaFold2 on 5SBV-model-3 (a), 7B5U-model-5 (b), T1031-model-4 (c), T1039-model-1 (d), T1029-model-3 (e) and T1033-model-4 (f). In each line, the first subgraph (gray) represents the native structure after alignment. The second to fourth subgraphs all represent the decoy structure, and the colors on the structure represent the score distribution of different evaluation methods (*LDDT*, score of QATEN, *pLDDT* of AlphaFold2), and the mean value of the score are shown under structures. The range of color (red-orange-yellow-green-cyan-blue-purple) is correlated with the range of scores (1-0).

As can be seen from Fig.7a and Fig.7b, AlphaFold2 predicts 5SBV-model-3 and 7B5U-model-5 quite well but AlphaFold2 is so confident that overestimates these decoys. The confidence predictor ignores some domains, resulting in low scores under 𝒫 (0.5938 and 0.6328). QATEN gives lower evaluation scores than *LDDT*, but achieves higher 𝒫 (0.7178 and 0.7555) than that of *pLDDT*. It suggests QATEN could indicate the domain need improvement in the decoys where AlphaFold2 performs well but overestimates. As shown in Fig.7c and Fig.7d, AlphaFold2 predicts T1031 and T1039 well but the confidence predictor in AlphaFold2 provides poor evaluation. 𝒫 of *pLDDT* are 0.3895 and 0.5996. QATEN provides closer assessment scores to *LDDT*, and achieves higher 𝒫 as 0.6 and 0.7281. Fig.7e and Fig.7f show two decoys relatively poorly predicted and worse evaluated (T1029-model-3 and T1033-model-4), QATEN performs much better on those decoys.

We also generate comparison graphs for all decoys (648) where 𝒫 by QATEN is higher than that by confidence predictor in AlphaFold2 as shown in our network (http://www.csbio.sjtu.edu.cn/bioinf/QATEN/). The accurate quantitative QA scores enable users to not only estimate the prediction quality quickly but also identify the regions requiring further consideration and refinement.

### 3.6 Comparison of QATEN with QA module in RosettaFold

QA module in RosettaFold uses predicted *C*_*α*_ -*RMSD* (*pRMSD*) to locally evaluate its predicted decoy structure, thus the correlation coefficients are obtained by calculating *pRMSD* and *RMSD*. We compare QATEN and the QA module in RosettaFold separately on CASP14_rosettafold and RCSB_rosettafold datasets. For 120 decoys in CASP14_rosettafold, the average 𝒫 of local assessment is 0.664 in QATEN and 0.6138 in QA module in RosettaFold. For 1230 decoys in RCSB_rosettafold, the average 𝒫 of local assessment is 0.6838 in QATEN and 0.6066 in QA module in RosettaFold. QATEN provides a more precise assessment of both CASP14_rosettafold and RCSB_rosettafold datasets.

Likewise, we also define a hit decoy means 𝒫 of a model on current decoy is higher than 0.5 and a hit protein means 𝒫 of a model on all 5 decoys of current protein are higher than 0.5. Table 4 shows the number of hit decoys and hit proteins by QATEN and QA module in RosettaFold. It is worth noting that the decoys used in these evaluations are all predicted by RosettaFold itself, the advantage of QATEN indicates possible support for the structural prediction of RosettaFold by QATEN.

**Table 4.** Comparison between QATEN and the QA module in RosettaFold

| Dataset | Method | $N_{heat_{decoy}}^a$ | $N_{heat_{protein}}^b$ |
| --- | --- | --- | --- |
| RCSB_rosettafold | **QATEN** | **1192** | **228** |
| | $pRMSD$ | 823 | 128 |
| CASP14_rosettafold | **QATEN** | **102** | **17** |
| | $pRMSD$ | 90 | 14 |

<sup>a</sup>: hit number of decoys; <sup>b</sup>: hit number of proteins.

We also merge CASP14_rosettafold and RCSB_rosettafold and further explore from the perspective of MSA depth (the MSA is searched and generated by RosettaFold) and secondary structure. MSA depth in Fig.8 is generally higher than that in Fig.5 because Rosetta ignores redundancy when searching for homologous information. As shown in Fig.8, QATEN performs better on decoys with different MSA depths and almost hits all the decoys (97.2%, 1165 decoys in total) with MSA depths from 2000 to 20000.

**Fig.8.**
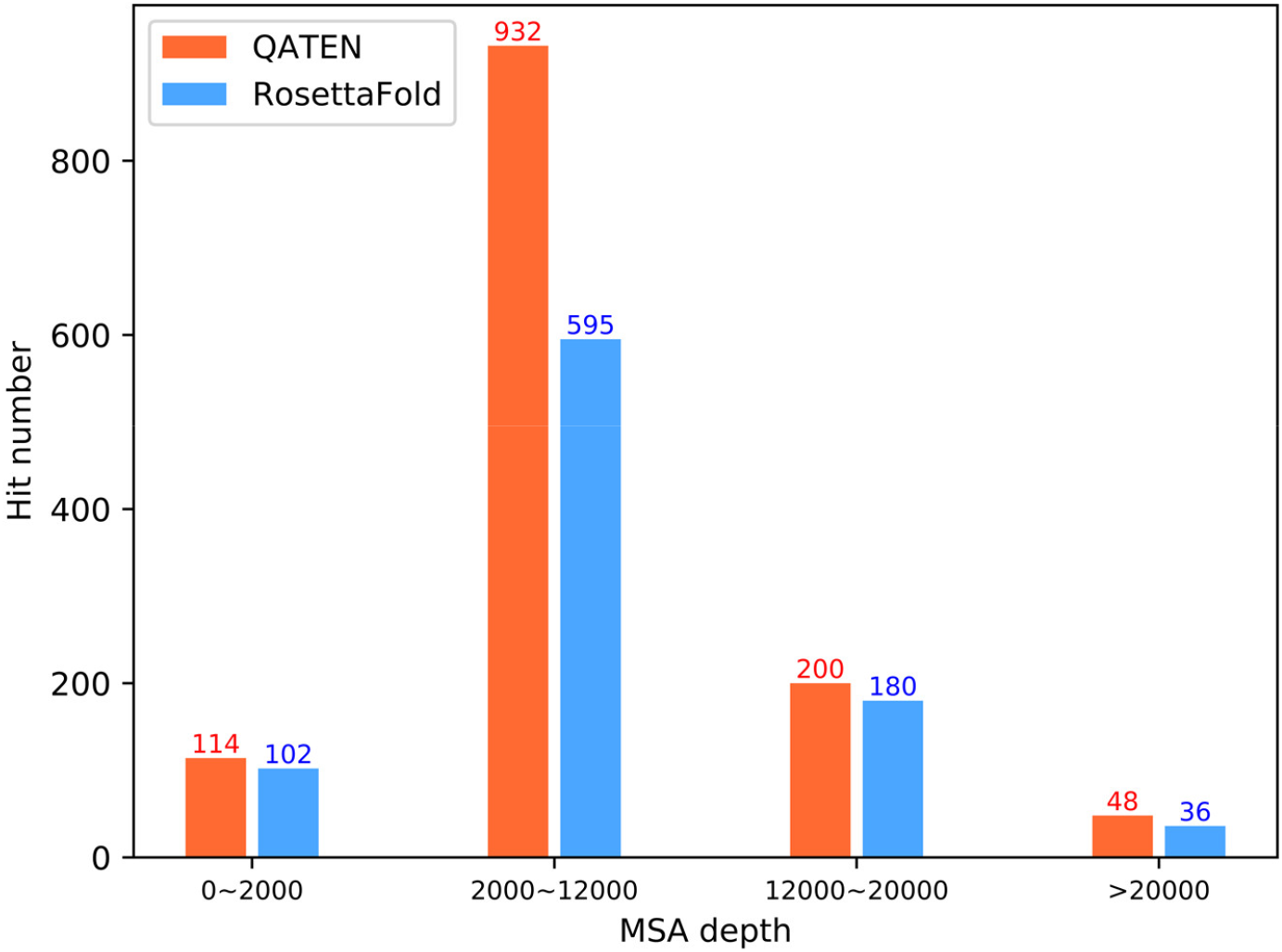
Distribution of the hit decoys according to MSA depth. QATEN performs better on decoys with different MSA depths and almost hit all decoys (97.2%) with MSA depths from 2000 to 20000.

We find 416 decoys only hit by QATEN and 35 decoys only hit by QA module in RosettaFold, and count the distribution of secondary structure elements using DSSP [37] software. As shown in Fig.9, the distribution of secondary structure elements in decoys only hit by (a) QATEN or (b) RosettaFold. Additionally, the distribution of secondary structure elements in all decoys is also shown in (c). The distribution of secondary structure elements in two sets mainly concentrated on Strand and Helix. Compared to general decoy in CASP14_alphafold and RCSB_alphafold datasets, decoys only hit by QATEN do not show a significant difference. And a slightly less helix (-2.9%) could be observed on those decoys only hit by QA module in RosettaFold. These results indicate that QATEN could be used as an extra QA tool for the decoys generated by RosettaFold.

**Fig.9.**
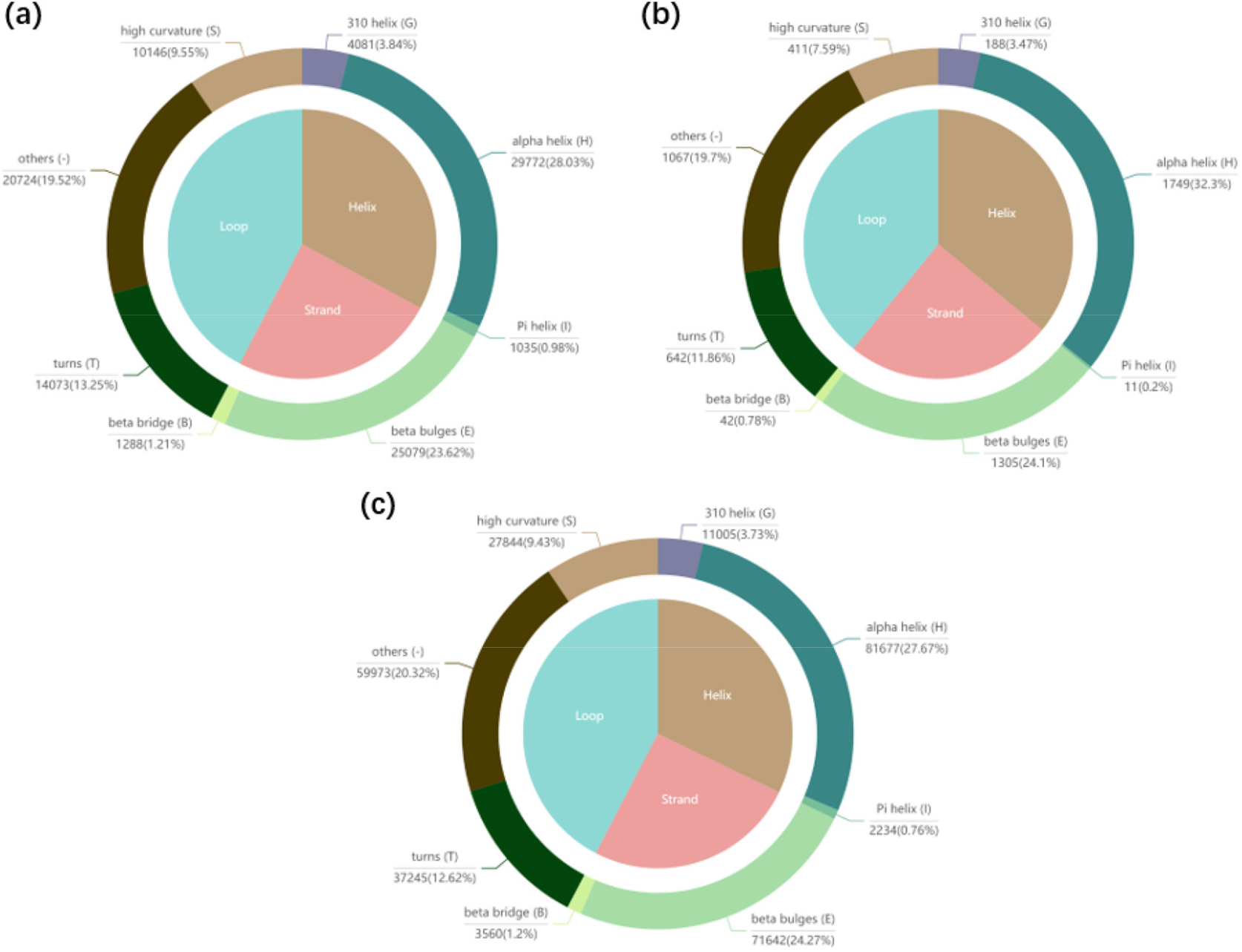
Distribution of secondary structure elements in decoys only hit by (a) QATEN or (b) QA module in RosettaFold. Additionally, the distribution of secondary structure elements in all decoys is also shown in (c).

In Fig.10, we make six cases comparison between the QATEN and the RosettaFold assessment *pRMSD*. The structural prediction module in RosettaFold performs quite well for the model has been improved before we download. Each graph represents the decoy sample predicted by RosettaFold, and the six graphs represent 7FH7-model-1, 7PXY-model-4, T1025-model-4, T1030-model-5, T1042-model-4 and T1079-model-4 respectively. The first subgraph (gray) represents the native structure after alignment. The second to fifth subgraphs all represent the decoy structure, and the colors on the structure represent the score distribution of different evaluation methods (*LDDT*, score of QATEN, *RMSD, pRMSD* of RosettaFold), and the mean value of the score is shown under structures. Particularly, *RMSD* and *pRMSD* are back-normalized to be between 0 and 1 for ease of comparison. The range of color (red-orange-yellow-green-cyan-blue-purple) is correlated with the range of scores (1-0).

**Fig.10.**
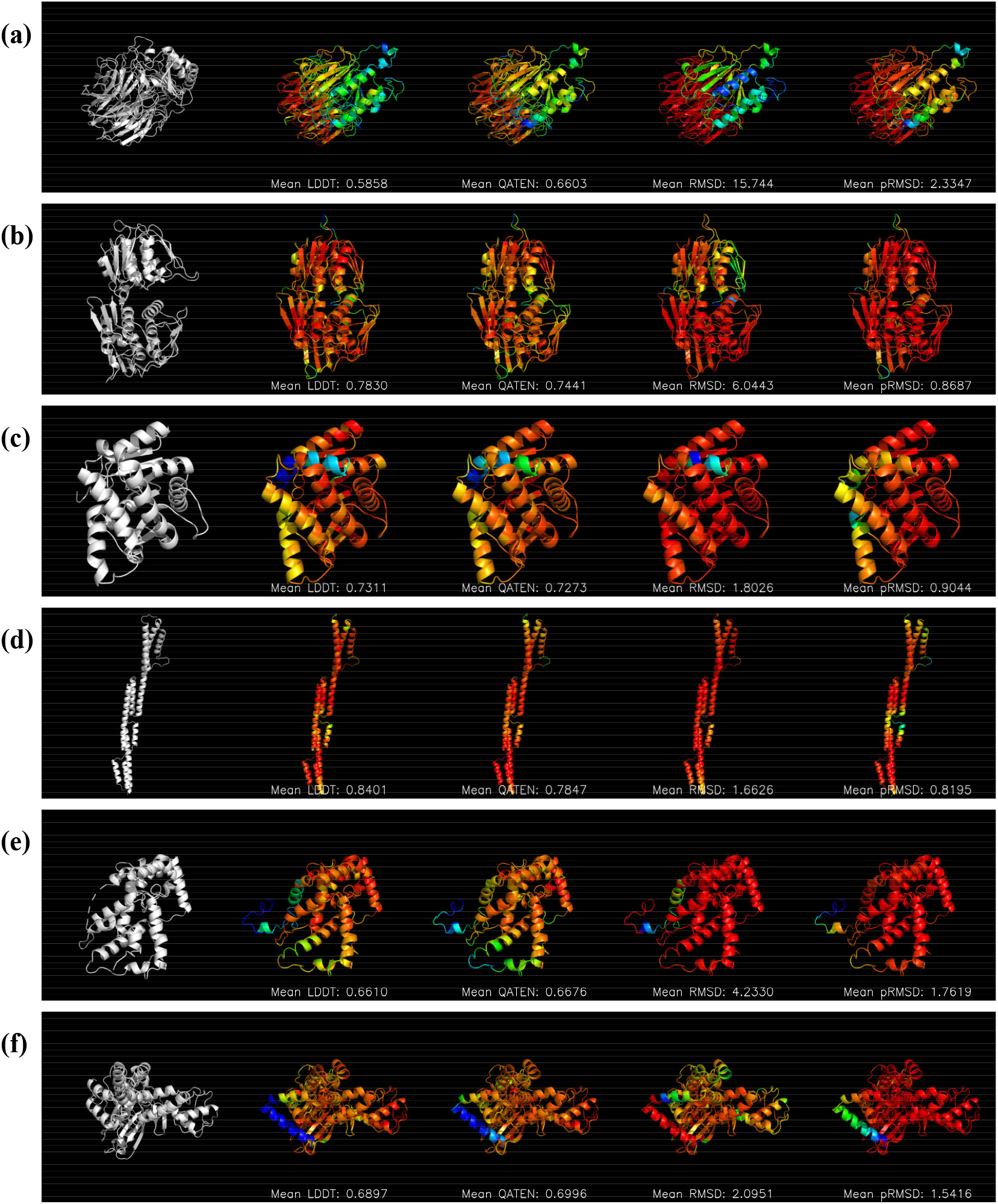
Comparison of QATEN and *pRMSD* of RosettaFold on 7FH7-model-1 (a), 7PXY-model-4 (b), T1025-model-4 (c), T1030-model-5 (d), T1042-model-4 (e) and T1079-model-4 (f). In each line, the first subgraph (gray) represents the native structure after alignment. The second to fifth subgraphs all represent the decoy structure, and the colors on the structure represent the score distribution of different evaluation methods (*LDDT*, score of QATEN, *RMSD, pRMSD* of RosettaFold), and the mean value of the score is shown under structures. Particularly, *RMSD* and *pRMSD* are back-normalized to be between 0 and 1 for ease of comparison. The range of color (red-orange-yellow-green-cyan-blue-purple) is correlated with the range of scores (1-0).

As can be seen from Fig.10c and Fig.10d, RosettaFold predicts these two decoys well but QATEN can provide a more precise evaluation (𝒫 as 0.9035 and 0.7444) than the assessment module of RosettaFold (𝒫 as 0.39 and 0.5366) at decoys with simple structures. Fig.10b and Fig.10f show QA module in RosettaFold is so confident that it overestimates and ignores some domains of two decoys that need improvement (𝒫 as 0.202 and 0.3681). QATEN precisely identifies these domains for improvement and achieves higher 𝒫 (0.7541 and 0.7569). Fig.10a and Fig.10d show two decoys RosettaFold relatively poorly predicted. QATEN provides more accurate evaluations (𝒫 as 0.8682 and 0.7444) than QA module in RosettaFold does (𝒫 as 0.5425 and 0.5366).

We also generate comparison graphs for all decoys (862) where 𝒫 by QATEN is higher than that by QA module in RosettaFold, which could be found in our network (http://www.csbio.sjtu.edu.cn/bioinf/QATEN/).

### 3.7 Computational analysis

As mentioned earlier, QATEN only uses atomic type and topology features as its inputs. At the same time, the trainable parameters of the entire network are ∼80k. We randomly select 1000 decoy structures for prediction using QATEN, ProteinGCN, DeepAccNet, DeepAccNet-bert, and ZoomQA respectively. All prediction processes are based on a single thread in a 2.4GHz Intel CPU and are timed separately. Table 5 shows the speed comparison between QATEN and the software we downloaded running locally. As can be seen from Table 5, QATEN spends the least time predicting 1000 decoys in total. ProteinGCN also costs relatively little time because it uses fewer original features than that of other models. DeepAccNet and DeepAccNet-bert spend acceptable time when not searching and using multiple sequence alignment information. Although ZoomQA only uses SVM as training networks and does not use homology search and PSSM. It requires extracting features and making decisions for each amino acid in the protein chain, which greatly increases the time cost.

**Table 5.** Speed comparison between QATEN and baseline models on 1000 decoys.

| Methods | Cost time in total |
| --- | --- |
| QATEN | 12m 46s |
| ProteinGCN | 14m 32s |
| DeepAccNet | 31m 3s |
| DeepAccNet-bert | 33m 10s |
| ZoomQA | 4h 3m 0s |

## 4 Method

### 4.1 Feature extraction

A natural way to represent decoy structures is to model them as graphs, which have been shown to be effective in recent works [28, 29, 35, 38]. In this paper, we create a protein graph similar to ProteinGCN [28]. We consider the 50 nearest neighbor atoms in the three-dimensional space of each node atom and connect them to the node atom in the protein graph. In this protein graph, we use one-hot vector encoding as a feature to represent atomic types as node features; as for edge features, we specifically include the following three categories.

Edge distance: for every two atoms that will be connected in the protein graph, we calculate their relative Euclidean distance in the decoy. In this work, we use the Gaussian extension to generate a 40-d vector representation of the distance.

Edge coordinates: for each atom *B*, it can form a local reference frame with its pre-sequence atom *A* and post-sequence atom *C* in the amino acid sequence. Define ***c*** as the spatial vector formed by the three-dimensional coordinates of a pair of atoms. The three unit vectors ***e***_***x***_,*e*_*y*_,***e***_***z***_ are calculated as follows:

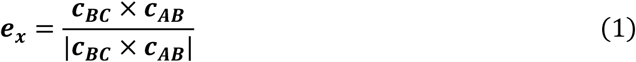

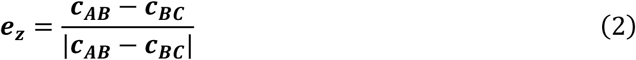

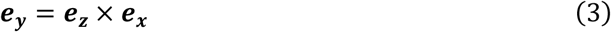

When considering the edge coordinates of any two atoms *B*_*i*_, *B*_*j*_, the three-dimensional space coordinates of *B*_*i*_ *and B*_*j*_ are replaced by the local reference system of each other for representation. Then the edge (*i, j*) in the protein graph obtained 2×3 features, respectively representing the projection of the spatial coordinates of the atom *B*_*i*_(*B*_*j*_) on the local coordinate system of the atom *B*_*j*_(*B*_*i*_).

Biological bond: a Boolean feature that is set to 1 if the connection between the two atoms is a chemical bond in an actual protein structure, and set to 0 otherwise.

### 4.2 Model architecture

QATEN consists of a five-layer main network and a downstream network based on graph pooling. The main network is built using an attention mechanism, and each layer includes a triple attention module like Transformer. The transformer has seen great success since it was proposed [39], and AlphaFold2 demonstrated it could be successfully applied to bioinformatics by designing the Evoformer module in CASP14 [6]. The main network obtains high-dimensional features and then connects to a pooling network, including two different graph pooling methods and fully connected layers. Formally, we will give the model formula below.

Given a protein graph **𝒢** *=* (**𝒜, 𝒱, 𝒰**), where ***a***_***ij***_ **∈ 𝒜** indicates whether the *ith* atom and *jth* atom are connected, ***v***_***i***_ **∈ 𝒱** indicates original node features of the *ith* atom, **µ**_***ij***_ **∈ 𝒰** indicates the edge features between the *ith* atom and *jth* atom.

#### High-dimensional feature extraction

The main network consists of 5-module frameworks with the same architecture and each module was designed like Evoformer. We use it on protein graph information rather than MSA information, and design a shortcut connection like ResNet to avoid overfitting. Fig.11 shows the architecture of one module. In Fig.11, *N* means the length of atoms, *M* means the number of neighbors, and *h* means the number of multi-heads. *c*_*a*_, *c*_*b*_ and *c*_*m*_ respectively represent the channels of node feature, edge feature, and reconstructed protein graph. The relationship between them is shown in the following formula.

**Fig.11.**
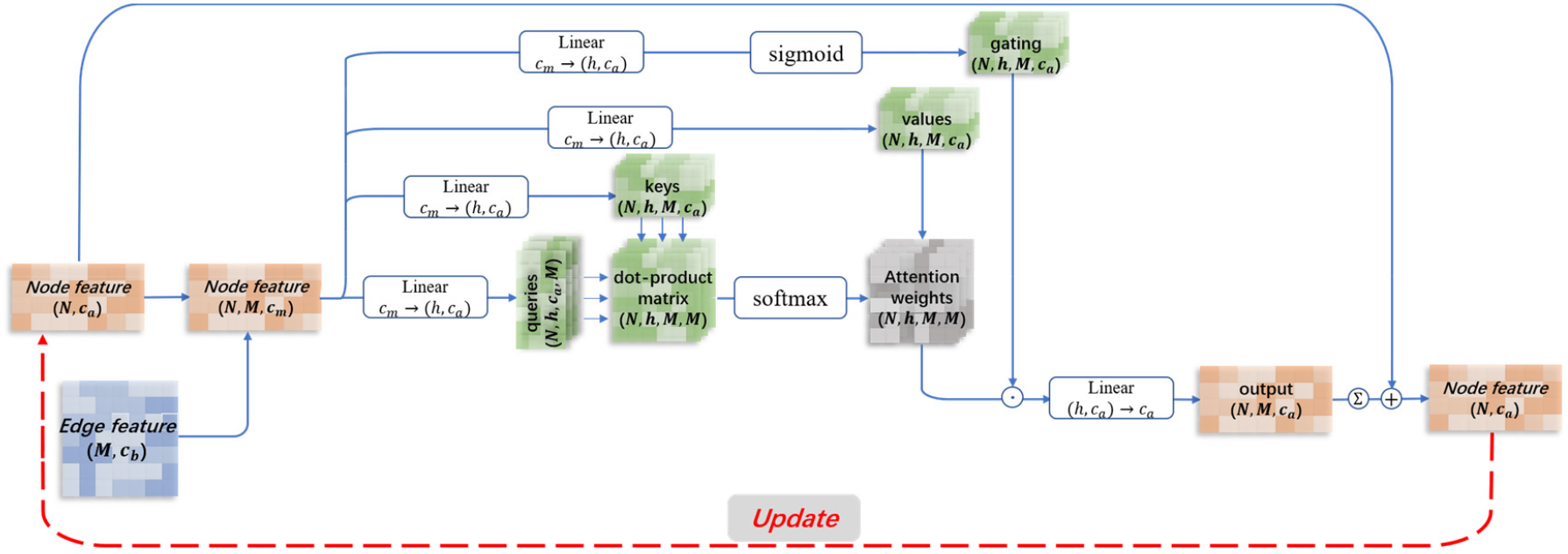
One module based on attention in GNNs. In each module, node features will be updated by the attention mechanism while edge features are unchanged.

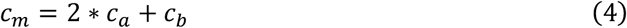

To further illustrate with formulas and for each module, the total calculation formula is as follows:

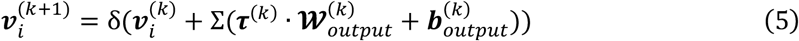

The superscript *k* in the formula means the *k*th module in the network, which can be 1, 2, 3, 4, 5. δ(·) represents RELU nonlinear activation function. Σ(·) represents summing tensors on the second dimension.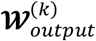 and 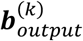 are the convolution weight matrix and convolution bias matrix in the fully connected layer that is finally used to capture the multi-head information. In addition, **τ**^(*k*)^ is the feature map calculating by attention mechanism:

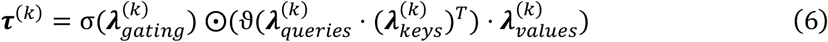

where 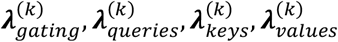 are four feature maps calculated respectively by four multi-head fully connected layers with reconstructed protein map ***Z***^(*k*)^ as input. ϑ(·) represents the softmax function, and σ(·) represents the sigmoid function. ⨀ means element-wise product.

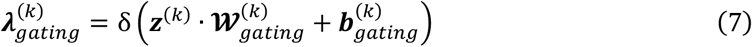

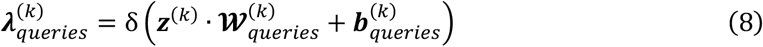

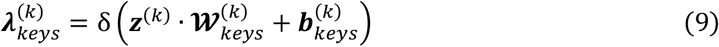

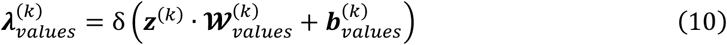

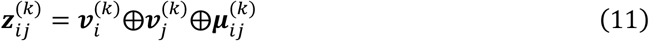

The ⨁ means concatenation between vectors.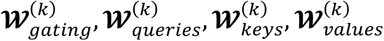 and 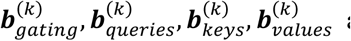 are the convolution weight matrix and convolution bias matrix in the fully connected layer of the *k*^*th*^ module. Respectively, the number of heads is set to 3 in our work.

#### Graph pooling

For the node embedding representation 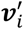 obtained by the forward calculation, the global and local scores are obtained by using the graph pooling technique.

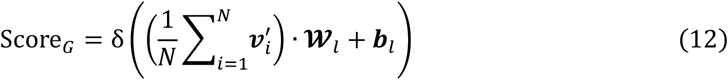

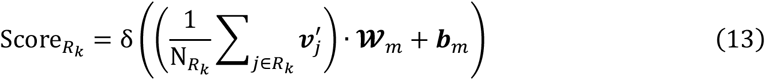

where Score_*G*_ and 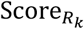 represent the final global score and the *k*^th^ amino acid’s local score. 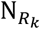 represents the set of atoms that make up the *k*^th^ amino acid, **𝒲**_*l*_, ***b***_*l*_,**𝒲** _*m*_, ***b***_*m*_ are the weight matrix and bias matrix in the global and local fully connected layer.

### 4.3 Loss function

We design a novel loss function for the training model to focus on high-precision decoys and the high-precision domain of decoys. Structural prediction models could not accurately predict unfamiliar protein sequence before CASP13. It was an easier challenge to select high-precision decoy structures at the protein level and identifies a domain that need improvement at the fragment level from different models at that time. Most QA models tend to design more complex networks and use more features, including MSA information, PSSM, and DSSP, to improve their results. General advances in structural prediction models have made it difficult for QA models to successfully rank high-precision decoys.

Researches have shown that it is relatively easy to identify decoys and domains that were incorrectly modeled. The current challenge is to accurately evaluate and rank the decoy structures with general or good accuracies and guide the direction of further local refinement. To this motivation, we design the loss function in formula (11). Both predictions and labels will have global and local parts, respectively. Assuming that a decoy consists of *L* amino acids, the final score obtained by the network is 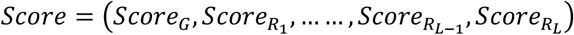, and the corresponding label is also 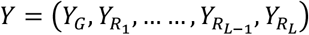, then the loss of the decoy is defined by the following formula:

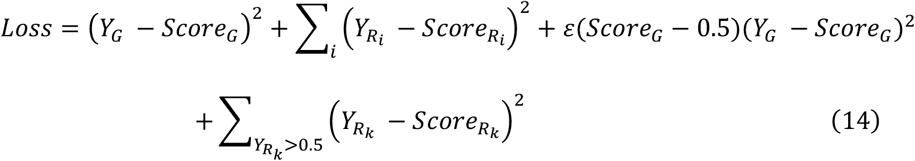

where ε(·) is the step function. The first and second terms of the formula (11) calculate the root mean square error (RMSE) between labels and predictions from global and local perspectives, like loss function in general QA models. The third term calculates RMSE between global labels and global predictions of high-accuracy decoys and the fourth term calculates RMSE between local labels and local predictions of high-accuracy domain in decoys.

Obviously, the loss function will focus on high-precision decoys and local segments in each decoy that are closer to the ground truth. It explains the success of QATEN on high-precision decoy datasets and decoys generated by AlphaFold2 and RosettaFold. The first and second terms of the loss function ensure that QATEN is also able to identify decoys modeled incorrectly. Besides, we do not enforce the weight of global labels in the loss function, resulting in QATEN being more successful in the local evaluation under *LDDT* analysis. The ablation experiments about loss function can be found in Supplementary information.

## 5 Conclusions

This paper presents a new method QATEN for protein quality assessment. QATEN uses graph neural networks (GNNs) and the attention mechanism to update node representation in protein graph from the decoys structural features, then concatenates and expands the predicted node features into a pooling network to obtain the final results. In order to get more accurate evaluations on high-accuracy decoys, we have designed new attention network architecture and loss functions. In our current implementation, QATEN does not make use of any sequence homologs and secondary structure features. Since only a few features and parameters are trained, QATEN is able to get the evaluation results very quickly. Evaluation of performance on multiple benchmark datasets shows that QATEN can derive accurate evaluations.

## Code availability

QATEN is available at: http://www.csbio.sjtu.edu.cn/bioinf/QATEN/, and source code is available at: https://github.com/CQ-zhang-2016/QATEN.

